# Transcriptional regulators FOXD1 and RBFOX2 contribute to metastatic capacity in *BAP1*^*mut*^ uveal melanoma

**DOI:** 10.1101/2025.04.14.648661

**Authors:** Q.C.C. van den Bosch, J.Q.N. Nguyen, Nikki Konemann, T.P.P. van den Bosch, R.M. Verdijk, E. Kilic, E. Brosens

**Affiliations:** Department of Ophthalmology, Erasmus MC University Medical Center Rotterdam, Rotterdam, The Netherlands; Clinical Genetics and Erasmus MC University Medical Center Rotterdam, Rotterdam, The Netherlands; Pathology Erasmus MC Cancer Institute, Erasmus MC University Medical Center Rotterdam, Rotterdam, The Netherlands

## Abstract

Uveal melanoma is the most prevalent primary intraocular cancer, with a significant metastatic risk. This risk is dependent on the genetic drivers. Secondary mutations in *EIF1AX, SF3B1* and *BAP1* correlate with clinical outcomes and are recognized for their distinct transcriptomic and epigenetic profiles. Previously, we identified 480 genes involved in the development of ocular melanocytes. Top ranking genes RBFOX2 and FOXD1 were significantly associated with BAP1 UM and were independently correlated to poor progression. However, it is uncertain whether either RBFOX2 or FOXD1 have biological contribution to disease progression or are solely indicative. This study investigates if and how these high-risk associated transcription regulators FOXD1 and RBFOX2 could influence tumor progression through knock-out and overexpression models. Our results indicate that loss of RBFOX2 affects cell morphology, attachment and proliferation, particularly in BAP1^neg^ cells. Additionally, both RBFOX2 and FOXD1 contribute to tumor growth and dissemination in zebrafish xenografts. Loss of either RBFOX2 or FOXD1 reduced tumor volume and cell dissemination, with the greatest effects seen in BAP1^neg^ cells. Overexpression models demonstrated different morphological and invasive behavior depending on the genetic background, suggesting complex roles in a context dependent fashion. Overexpression of RBFOX2 did not alter BAP1^pos^ cells yet made BAP1^neg^ cells more aggressive *in vitro* and *in vivo*. This study underscores the influence of RBFOX2 and FOXD1 as important factors for UM progression.

## Introduction

Uveal melanoma (UM) is the most common primary intraocular malignancy with a high risk of developing metastatic disease. Even after successful treatment of the primary tumor, up to 50% of patients will develop metastatic disease[1]. Clinical outcomes are associated with clinical, histological and genomic characteristics[2, 3]. Driver mutations in UM are typically found in *GNAQ[4]* or *GNA11* [5], and in a lower frequency in *PLCB4[6]* or *CYSLTR2[7]*. All driver mutations act in the MEK/ERK pathway, driving cell proliferation via YAP-signaling[2]. Driver mutations are found in benign ocular nevi [8, 9], illustrating the necessity of secondary mutations for malignant transformation. Secondary mutations in UM are typically identified in Eukaryotic Translation Initiation Factor 1A X-Linked (*EIF1AX)*, Splicing Factor 3b Subunit 1 (*SF3B1)* or deubiquitinase BRCA1-associated protein-1 *(BAP1)*, which are typically mutually exclusive. Secondary driver mutations strongly correlate to clinical outcome, stratifying patients into three distinct groups. UM patients harboring *EIF1AX*^*mut*^ have a low risk of developing metastasis, patients with a *SF3B1*^*mut*^ UM are considered intermediate risk, whereas patients with *BAP1*^*mut*^ UM have the highest risk of metastasis[10].

Despite clear correlations to metastatic risk in *BAP1*^*mut*^ patients, little is known about the exact regulatory mechanisms driving distant metastatic disease. However, gene expression analysis hints towards a dedifferentiated cell-state in *BAP1-*deficient cells [11]. A major component involved in gene regulation are transcription factors, which are known to determine how cells function and respond to the environment [12]. In cancer, transcriptional regulators are often mutated or dysregulated [13], resulting in transcriptional addiction [14]. Inhibition of transcription regulators yielded promising results in multiple cancers, especially when combined with another compound, e.g. a kinase inhibitor [13]. Regulatory elements causing loss of melanocytic cell identify in *BAP1*^*mut*^ UM remains largely unclear. Previously, we investigated transcription regulators expressed in early vertebrate melanocyte development and validated expression in human UM primary tumors [15]. The top-hit transcription regulators in this study were *FOXD1* and *RBFOX2*, which both correlated to poor prognosis and elevated in *BAP1*^*mut*^ UM. In this study, we aim to elucidate their mechanism and role in UM progression by generating stable knock-out and overexpression of either *FOXD1* or *RBFOX2* in *EIF1AX*^*mut*^*-* and *BAP1*^*mut*^ cell lines.

## Methods

### Generation of CRISPR-Cas9 Knock-out cell lines

Stable knock-out UM cell lines (92.1[16] and MP46[17]) were made using the PX458 plasmid (Addgene, Watertown, MA, US, #48138) containing guide RNA (gRNA) sequenced targeting *FOXD1* (*5’CATCGGACATCTCAGTGCTC*) or *RBFOX2* (5’*ACGGAAGTACGCAAGCCCAC*). UM Cell lines were transfected with Lipofectamine 2000 (ThermoFisher, Bleiswijk, Zuid-Holland, The Netherlands, Cat. no. 11668500) and FOXD1-or RBFOX2-PX458 plasmid. After 24 hours, cells were harvested and sorted based on GFP expression. A total of 5000 GFP+ cells were sequentially plated in a 10cm^2^ petri-dish and allowed to grow at 37°C with 5% CO_2_ in a humidified incubator until small colonies appeared (∼5-10 cells). Afterwards, colonies were manually picked and placed in a 96-wells plate and grown until enough cells were available for DNA isolation. DNA of selected clones was extracted by incubating cell pellets for 1 hour at 56°C in 10% Chelex solution supplemented with 1% Proteinase K. Supernatant was collected and used for PCR and Sanger sequencing. Amplification of *FOXD1* and *RBFOX2* was achieved using Platinum™ Taq DNA Polymerase (ThermoFisher) using the following primers: *FOXD1: 5’GGCAGCCGCAGTCGCA & 5’GCCCTCGTCGTCGTCCTCCT*; *RBFOX2: 5’GCATTCCATTTTACCTATGCAGTGAG & 5’ GTATTTCTAGGAGGAGGGAGGACAATG*. Sanger sequences were processed in TIDE[18]. Clones that lack any wild-type allele were used for further validation experiments.

### Generation of stable overexpression of RBFOX2 or FOXD1 in uveal melanoma cell lines

Stable overexpression of RBFOX1 or FOXD1 was achieved by lentiviral integration using adapted vectors. MP46 cells were unable to continue proliferation after puromycin selection, which made FACS essential to obtain a pure cell line. We utilized TFORF2004 (Addgene, 143330) as our backbone, as this plasmid already contains the FOXD1 cDNA and viral elements needed. We replaced the puromycin selection with a CMV-GFP from the pCMV-GFP plasmid (Addgene, 11153). This resulted in our pFOXD1-GFP plasmid (Supplementary Figure 4A), which we sequentially adapted by removing the FOXD1 cDNA and replacing it with RBFOX2 cDNA from pEGFP-RBFOX2 plasmid (Addgene, 63086) to obtain the pRBFOX2-GFP plasmid (Supplementary Figure 4B).

To obtain virus, HEK293T cells were seeded to obtain 80% confluence and transfected with pMDLg/pRRE (Addgene, 12251), pRSV-Rev (Addgene, 11253) pMD2.G (Addgene, 12259). After 4 hours, transfection medium was replaced by DMEM with 10% FCS and 1% NEAA (ThermoFisher, 111400500). Virus was isolated after 48 hours. Cell debris was removed using a 45um filter (Merck, SLHAR33SS). Isolated virus was directly used for transduction where a total of 1mL of virus containing medium with 8μg/mL polybrene was added to a 12-well containing either 92.1 or MP46 cells. After 24 hours, virus containing medium was removed and cells were allowed to expand for 1-2 weeks. Sequentially, GFP-positive cells were sorted using FACS to obtain a GFP-positive cell culture.

### Cellular architecture and attachment analysis

A total of 2.5×10^5^ cells were seeded in 48-wells plate and imaged on a Olympus IX-70 (Olympus Corporation, Tokyo, Japan) at the same coordinate at 1, 2, 3, 4, 6, 20 and 48 hours post-seeding. A total of 5 wells per cell line was investigated. Obtained images were stacked and analyzed in FIJI to manually count the number of attached and unattached cells. Sequentially, 2 additional images per well were obtained at 48 hours post-seeding in order to analyze cell shape of 40 attached cells per cell line. Cell detection was done manually in FIJI to measure cell perimeter, Feret’s diameter and circularity. Obtained values were processed in GraphPad V9 (GraphPad Software, Boston, MA, US) using Mann-Whitney T-Test comparing knock-out cells to WT-condition.

### Quantitative PCR

RNA was extracted from UM cell lines using QIAGEN miRNeasy Mini Kit (Qiagen, Hilden, Germany cat.no. 217084) according to manufactures protocol. Sequentially, cDNA was synthesized using iScript cDNA Synthesis kit (Bio-Rad) and used as template for quantitative PCR. Amplification of target genes and house-keeping gene was achieved using the following primers: *FOXD1: 5’CCCTGAGCACTGAGATGTCC & 5’CTCCTCCTCCTCCAGATCCT; RBFOX2: 5’GAGCAGAGCAGCAACTCACC & 5’GCCAAACTGCCCAAACATC; CHMP2A: 5’ AGAACCAGAGGGCCCTGAAC & 5’GAACAGCATCCATCTGGCCT*. Samples were amplified in a Bio-Rad CFX96 Touch Real-Time PCR Detection System (Bio-Rad Laboratory B.V.,Veenedaal, The Netherlands); Ct-values were extracted and analyzed using the ΔΔCt-method.

### Immunocytochemistry

UM cells were grown under standard conditions and harvested for immunocytochemistry (ICC). A total of 5000-7000 cells were used to generate a cytospin slide for stainings. ICC was carried out with an automated, validated and accredited staining system (Ventana Benchmark Discovery, Ventana Medical Systems, Tucson, AZ, USA). Following heat-induced antigen retrieval (CC1 for 4 min), cytospins were incubated with RBFOX2 antibody (1:800 dilution, ThermoFisher, PA5-52268) or FOXD1 (1:800 dilution, ThermoFisher, PA5-35145) for 32 minutes at 37°C. Secondary anti-rabbit-HRP was sequentially incubated for 20 minutes followed by thymidine based FAM detection for 4 minutes. Slides were covered with VectaShield (Vector Laboratories,Newark, CA, US, cat.no.H-1000-10) containing DAPI (1:1000 dilution). Fluorescent images were processed in QuPath V0.5 for protein quantifications. DAPI signal was used to automatically identify cell nuclei and FAM-signal was measured per nuclei to obtain RBFOX2 protein levels per UM Cell line. Fluorescent unit values were then plotted using GraphPad V9 and statistically analyzed using ANOVA testing. Fluorescent signal below arbitrary unit 200 were considered as background.

### Immunohistochemistry

Immunohistochemistry was performed with an automated, validated, and accredited staining system (Ventana Benchmark ULTRA, Ventana Medical Systems, Tucson, AZ, USA) using an ultraview Universal Alkaline Phosphatase Red Detection Kit (Ventana reference no. 760-501). Following deparaffinization and heat-induced antigen retrieval (CC1 for 64 min), the tissue samples were incubated for 32 min with the RBFOX2 antibody (1:800 dilution, ThermoFisher, PA5-52268). Counterstain was done by hematoxylin II stain for 12 min and a blue coloring reagent for 8 min according to the manufacturer’s instructions (Ventana Benchmark ULTRA, ThermoFisher, PA5-52268). Slides were scored by an ophthalmic-pathologist into an intensity matrix from 1+ up to 3+, representing low, intermediate and high expression. Corresponding intensity scores were plotted against clinical outcome in GraphPad Prism V9.

### Wound healing assay

*In vitro* migration capacity was assessed using Culture-Insert 3 well dish (Ibidi,Gräfelfing, Germany, cat.no. 80366) by seeding UM cell lines on the outer sides while seeding fibroblasts (GM22136, obtained from American Type Culture Collection, Manassas, VA, USA) in the middle well. A day after seeding, culture inserts were removed, and cells were washed twice with PBS. Sequentially, RPMI 1640 medium with Pen/Strep without serum was added to minimize proliferation during imaging. Cells were imaged with a Nikon Wide Field microscope every 10 minutes for a total time of 24 hours under standardized conditions (humidified, 5% CO_2_, 37°C). A total of 16 cells were analyzed per cell line in FIJI using the manual tracking plugin to obtain total migration distances. Data was plotted in GraphPad V9 and statistically tested using Mann-Whitney ANOVA.

### Cell Proliferation assay

Cell proliferation was analyzed using the Opera Phenix high content screening system (Perkinelmer, Shelton, CT, US). A total of 5000 cells were seeded and imaged every 12 hours for 3 days in total. Each cell lines contained 5 replicates. Growth was assessed based on the global surface coverage area. Data was imported into GraphPad V9 and transformed using Y=Y/K, where K= the average of time point 0 hours of 5 independent measurements. After transformation, nonlinear regression using exponential (Malthusian) growth was performed to obtain doubling times. Additionally, the transformed data was subjected to 2way ANOVA testing for statistical differences between cell lines.

### Zebrafish xenograft models

Wild-type AB zebrafish were maintained under standard conditions with an 14hr light and 10hr dark cycle. In this study, only larval zebrafish (no older than 120 hour post fertilization) were used. Animal experiments were approved by the Animal Experimentation Committee at Erasmus MC, Rotterdam. Zebrafish embryos were raised in E3 medium supplemented with 0.003% phenylthiourea in a petri dish at 28°C. At 24 hours post fertilization, medium was refreshed following dechorionation of larvae. At 48 hours post fertilization zebrafish larvae were anesthetized with 0.016% Tricane and used for injections. A total of 2,5×10^6^ cells were harvested and stained with 2,5μM CellTracker CM-Dil dye (Thermofisher, Bleiswijk, Zuid-Holland, The Netherlands, cat.no. C7000) for 5 minutes at 37°C and then for an additional 15 minutes at 4°C. After staining, CM-Dil dye was removed by centrifugation. The cells were then washed with PBS and suspended in 2% PVP-40/PBS. A total of ∼200-300 cells were injected in the perivitelline space. Successfully injected larvae were selected 1 hour post injection under a fluorescent stereomicroscope and placed in E3 medium with PTU and raised at 34°C. At 3 days post injection xenograft larvae were anesthetized and embedded in 1% low-melting agarose for live-cell imaging.

### Confocal microscopy of zebrafish xenograft

Zebrafish xenograft larvae were imaged using a Leica SP5 (Leica, Wetzlar, Germany) under standard conditions (561nm, 35% laser power with additional bright field image). Tile-scans of 3 images were generated to obtain full-body length images for analysis. Identification of disseminated cells, distance of dissemination and total tumor volume were calculated in FIJI using publicly available scripts described in previous work[19]. Obtained values were then processed in GraphPad Prism V9. Comparison of amount of detected spot, distance of dissemination and tumor volume between the different cell lines were statistically tested using Mann-Whitney T-Test to compare knock-out cells with wild-type condition.

## Results

### High RBFOX2 protein correlates to poor patient survival

Increased RNA expression of *RBFOX2* was previously described to correlate to overall survival [15]. To validate this finding, a total of 57 patient primary tumor FFPE slides were stained using immunohistochemistry, which illustrated a range of protein expression (Figure 1A). Staining patterns were divided into 3 classes: low expression (1+), intermediate expression (2+) and high expression (3+). Similar to the RNA expression levels, increased protein expression of RBFOX2 is significantly correlated to poor prognosis. In line with this transcription factor, we previously reported expression of FOXD1 to be correlated to poor prognosis on both RNA and protein levels [15]. Therefore, we continued to investigate both transcription factors to understand their contribution to tumor progression using functional assays.

**Figure 1:**
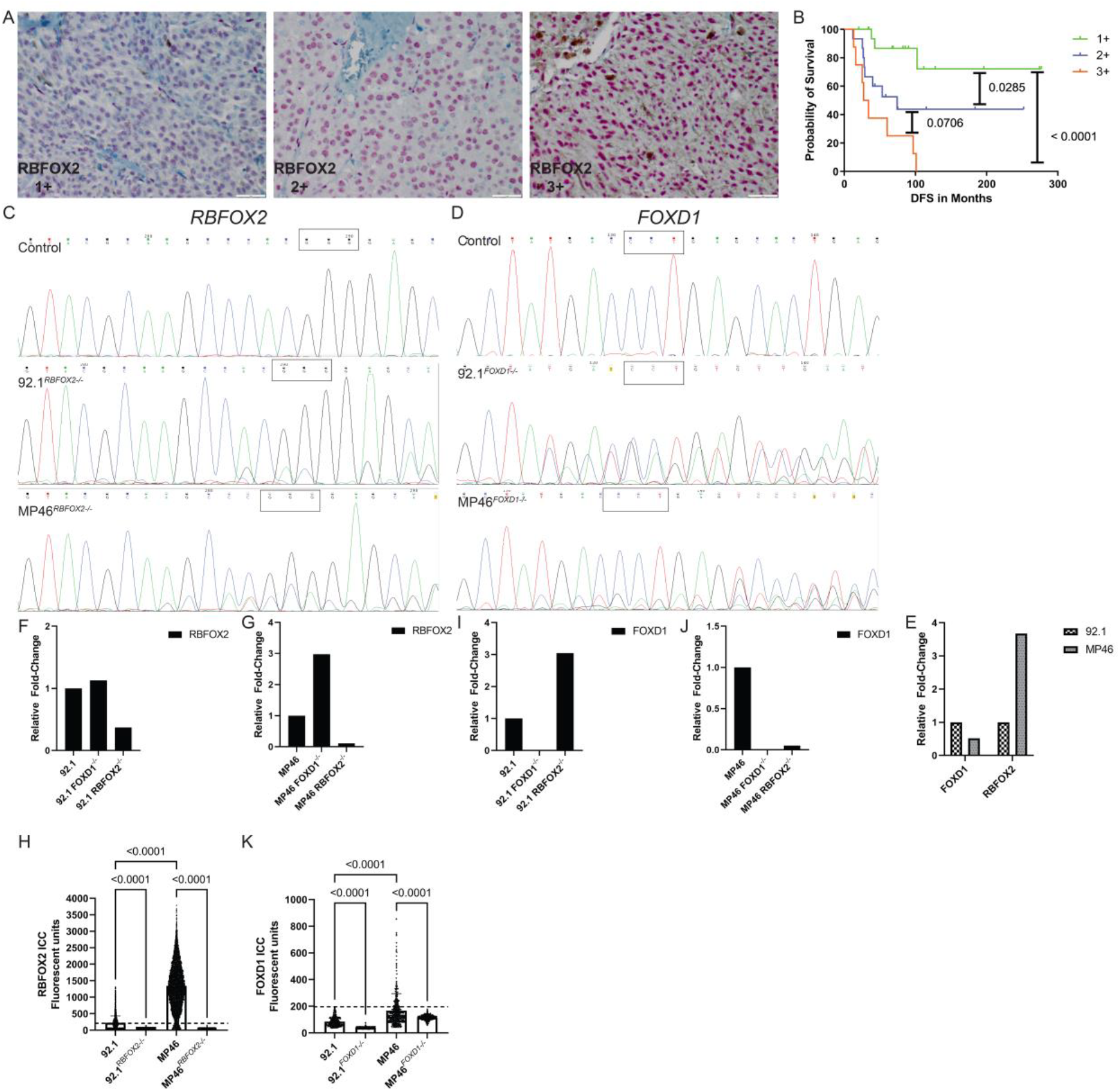
Validation of RBFOX2 protein expression and generation of Knock-out cell lines. A) Immunohistochemistry of UM patient samples illustrating low-(1+), intermediate (2+) and high (3+) expression of RBFOX2. B) Kaplan-Meier survival plot based on overall survival shows stratification of patients based on RBFOX2 expression. C) Sanger sequences of 92.1 cells in control condition, FOXD1 and RBFOX2 knock-out. D) Sanger sequences of MP46 cells in control condition, FOXD1 and RBFOX2 knock-out. F, G) RNA expression of RBFOX2 in WT and knock-out cells of 92.1 and MP46. I,J) RNA expression of FOXD1 in WT and knock-out cells of 92.1 and MP46. E) RNA expression comparison between WT 92.1 and WT MP46. H) Immunocytochemistry of RBFOX2 in WT and RBFOX2^-/-^ cells. K) Immunocytochemistry of FOXD1 in WT and FOXD1^-/-^ cells.

### Validation of stable uveal melanoma knock-out cell lines

To investigate how *RBFOX2* and *FOXD1* contribute to disease progression, we generated stable knock-out cell lines in BAP1-positive (92.1) and BAP1-negative (MP46) UM cell lines. Transient expression of a CRISPR-Cas9 plasmid containing one guideRNAs targeting *RBFOX2* or *FOXD1* allowed for generation of stable knock-out cell lines. Genomic editing differed per cell line, importantly clones used in this study show full loss of the wild-type allele (Figure 1C, 1D & Supplementary Figure 1A, 1B). RNA analysis demonstrated strong loss of *RBFOX2* mRNA transcripts in the *RBFOX2*^*-/-*^ cells (Figure 1F, 1G); and a complete loss of protein (Figure 1H, Supplementary figure 1C). RNA analysis of *FOXD1*^*-/-*^ cells illustrated a complete lack of mRNA transcripts (Figure 1I, 1J) and complete loss of protein (Figure 1K, Supplementary figure 1D). Due to strong reduced RNA transcripts and complete loss of protein, we considered these as stable knock-out UM cell lines and used these clones for this study. Additionally, based on the protein level, MP46 (BAP1^pos^) yields the highest expression of both RBFOX2 and FOXD1(Figure 1H, 1K).

### Loss of RBFOX2 alters morphology and inhibits cell attachment, invasion and proliferation

Morphological assessment of cellular shape illustrated that loss of RBFOX2 alters cellular architecture, whereas FOXD1 did not (Figure 2A-I). Quantifying cell perimeter and diameter demonstrated that *RBFOX2*^*-/-*^ UM cells are smaller with the largest effect seen in BAP1^neg^ cells (Figure 2B, 2D, 2G, 2I). Noteworthy, MP46^*RBFOX2-/-*^ cells are significantly more circular compared to MP46 cells (Figure 2H); an effect not seen in BAP1^pos^ cells (92.1, Figure 2C). During culture, it was evident *RBFOX2*^*-/-*^ had difficulties with attaching to the culture flask. Therefore, a time-based experiment was carried out to quantify attachment abilities of knock-out cell lines (Supplementary figure 2). Knock-out of *RBFOX2* resulted in slower attachment in vitro, where the biggest difference was seen in BAP1^neg^ cells (Figure 2E, 2J, supplementary figure 2). Loss of *FOXD1* have mixed results, which delayed attachment in BAP1^pos^ cells until 6 hours post seeding before it came back to levels comparable to WT condition (Figure 2E, supplementary figure 2). Loss of RBFOX2 in BAP1^neg^ cells on the other hand demonstrated an increase of attachment speed (Figure 2J, Supplementary figure 2). Finally, cell proliferation assays illustrated no alterations in doubling time after loss of either FOXD1 or RBFOX2 in BAP1^pos^ cells (Figure 2M, best fitted doubling times were 92.1 = 32.48H, 92.1^RBFOX2-/-^=34.38H and 92.1^FOXD1-/-^=34.19H). Interestingly, loss of RBFOX2 in BAP1^neg^ cells increased doubling time significantly while loss of FOXD1 did not show differences (Figure 2N, best fitted doubling times were MP46=69.79H, MP46^RBFOX2-/-^=53.99H and MP46^FOXD1-/-^ =59.51H).

**Figure 2:**
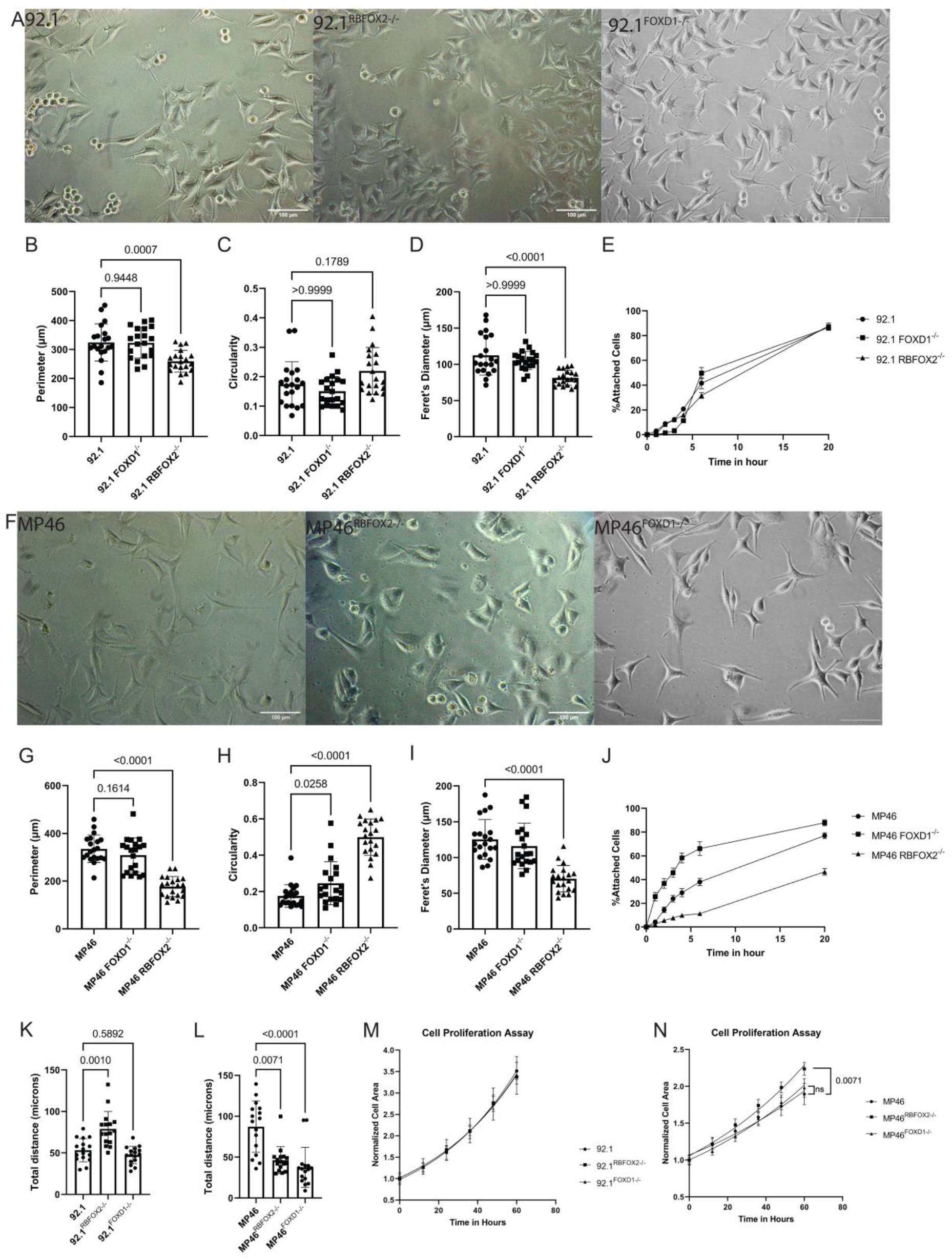
Morphological alterations and in vitro cell behavior after loss of FOXD1 or RBFOX2. A) Representative images of 92.1, 92.1^RBFOX2-/-^ and 92.1^FOXD1-/-^. B-D) Morphometric analysis of 92.1, 92.1^RBFOX2-/-^ and 92.1^FOXD1-/-^ on cell perimeter, circularity and Feret’s diameter, respectively. E) Cell attachment assay of 92.1, 92.1^RBFOX2-/-^ and 92.1^FOXD1-/-^ over the course of 20 hours. F) Representative images of MP46, MP46^RBFOX2-/-^ and MP46^FOXD1-/-^. G-I) Morphometric analysis of MP46, MP46^RBFOX2-/-^ and MP46^FOXD1-/-^ on cell perimeter, circularity and Feret’s diameter, respectively. J) Cell attachment assay of MP46, MP46^RBFOX2-/-^ and MP46^FOXD1-/-^ over the course of 20 hours. K-L) Total migration distance of 92.1, 92.1^RBFOX2-/-^,92.1^FOXD1-/-^, MP46, MP46^RBFOX2-/-^ and MP46^FOXD1-/-^ after 24 hours using the wound healing assay, respectively. M) Normalized cell surface area demonstrating cell growth of 92.1, 92.1^RBFOX2-/-^ and 92.1^FOXD1-/-.^ N) Normalized cell surface area demonstrating cell growth of MP46, MP46^RBFOX2-/-^ and MP46^FOXD1-/-^.

### Loss of *FOXD1* and *RBFOX2* in BAP1^neg^ UM cells inhibits tumor growth and cell dissemination abilities *in vivo*

Due to the significant alteration seen *in vitro*, zebrafish xenograft models were established to assess cellular behavior *in vivo*. A representation of the different xenograft models made using WT and knock-out cells are shown in Figure 3A-F. Quantification of xenograft models at 3 days post injection (3dpi) demonstrated loss of *RBFOX2* or *FOXD1* in BAP1^pos^ cells resulted in less tumor volume (Figure 3G). Loss of *RBFOX2* also reduced the number of disseminated cells and their dissemination distance (Figure 3H-I); which was not seen in *FOXD1*^*-/-*^ BAP1^pos^ cells. On the other hand, loss of *RBFOX2* or *FOXD1* showed a much stronger inhibitory effect in BAP1^neg^ cells. Tumor volume analysis of BAP1^neg^ cells showed significantly lower tumor burden in knock-out cells (Figure 3K). Cell dissemination distance and number of detected spots was also significantly reduced in both *RBFOX2*^*-/-*^ and *FOXD1*^*-/-*^ BAP1^neg^ cells (Figure 3L, M). Additionally, we investigated the number of large disseminated cell clusters with a size larger than 200 voxels. With the magnification used in this system, 10-20 voxels are representative for a single cell, therefore a spot of >200 voxels contain at least 10 cells. This could represent potential metastatic sites where cells are proliferating. However, the number of large spots were unaffected in both BAP1^pos^ and BAP1^neg^ cells (Figure 3J, N).

**Figure 3:**
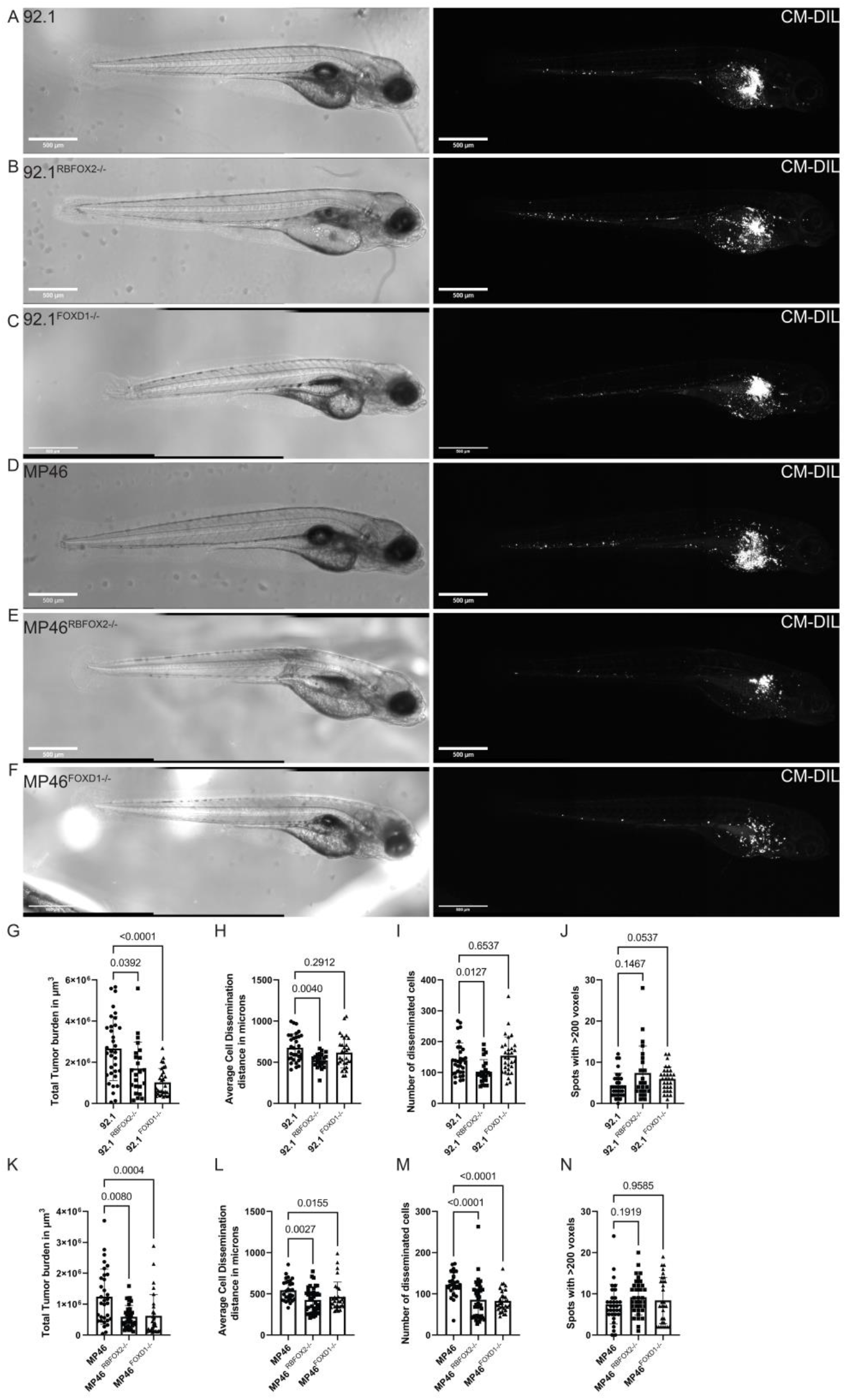
Overview of zebrafish larvae xenografts. A) 92.1-xenografts at 3 dpi. B) 92.1^RBFOX2-/-^-xenografts at 3dpi. C) 92.1^FOXD1-/-^-xenografts at 3dpi. D) MP46-xenografts at 3 dpi. E)MP46^RBFOX2-/-^-xenografts at 3dpi. F) MP46^FOXD1-/-^-xenografts at 3dpi. G) Tumor volume analysis of 92.1 WT and Knock-out cells in vivo. H) Cell dissemination distance analysis of 92.1 WT and Knock-out cells in vivo. I) Number of detected spots of 92.1 WT and Knock-out cells in vivo. J) Identification of large disseminated cell clusters with a larger then 200 voxel size in 92.1-based xenografts K) Tumor volume analysis of MP46 WT and Knock-out cells in vivo. L) Cell dissemination distance analysis of MP46 WT and Knock-out cells in vivo. M) Number of detected spots of MP46 WT and Knock-out cells in vivo. N) Identification of large disseminated cell clusters with a larger than 200 voxel size in MP46-based xenografts.

### In vitro characterization after overexpression of RBFOX2 and FOXD1 in uveal melanoma cell lines

Overexpression cell lines were generated using lentiviral integration of full-length cDNA of either *FOXD1* or *RBFOX2* in 92.1 and MP46 cells. After FACS of GFP+ cells, cells were validated for overexpression using Immunocytochemistry. All cell lines were successfully established (Figure 4 and Supplementary figure 5), with exception of FOXD1 overexpression in MP46. After multiple attempts, sorted MP46-FOXD1-GFP cells failed to proliferate and ultimately perished after 3 weeks of culture. Yet, 92.1-FOXD1-GFP cells were successfully established. This might suggest too high expression of FOXD1 can be detrimental in cells that naturally have elevated expression, although this expression is relatively low (Figure 1K). Following validation of protein overexpression (Figure 4A), morphometric and migration analysis followed as performed on knock-out cells. Interestingly, there were no significant changes in morphology besides a small increase in size of 92.1-RBFOX2-GFP cells (Supplementary Figure 6). Wound healing assays demonstrate an inhibited migratory capacity upon overexpression of FOXD1 in 92.1 cells, whereas overexpression of RBFOX2 did not (Figure 4C). However, overexpression of RBFOX2 in MP46 cells did significantly increase migratory distance compared to GFP control cells; suggesting RBFOX2 effect could be cell type specific (Figure 4D). This is further supported by cell proliferation differences between 92.1 and MP46 cells. Overexpression of FOXD1 or RBFOX2 severely reduced doubling times compared to GFP conditions in 92.1 cells (Figure 4E), whereas overexpression of RBFOX2 in MP46 cells improved doubling times significantly (Figure 4F).

**Figure 4:**
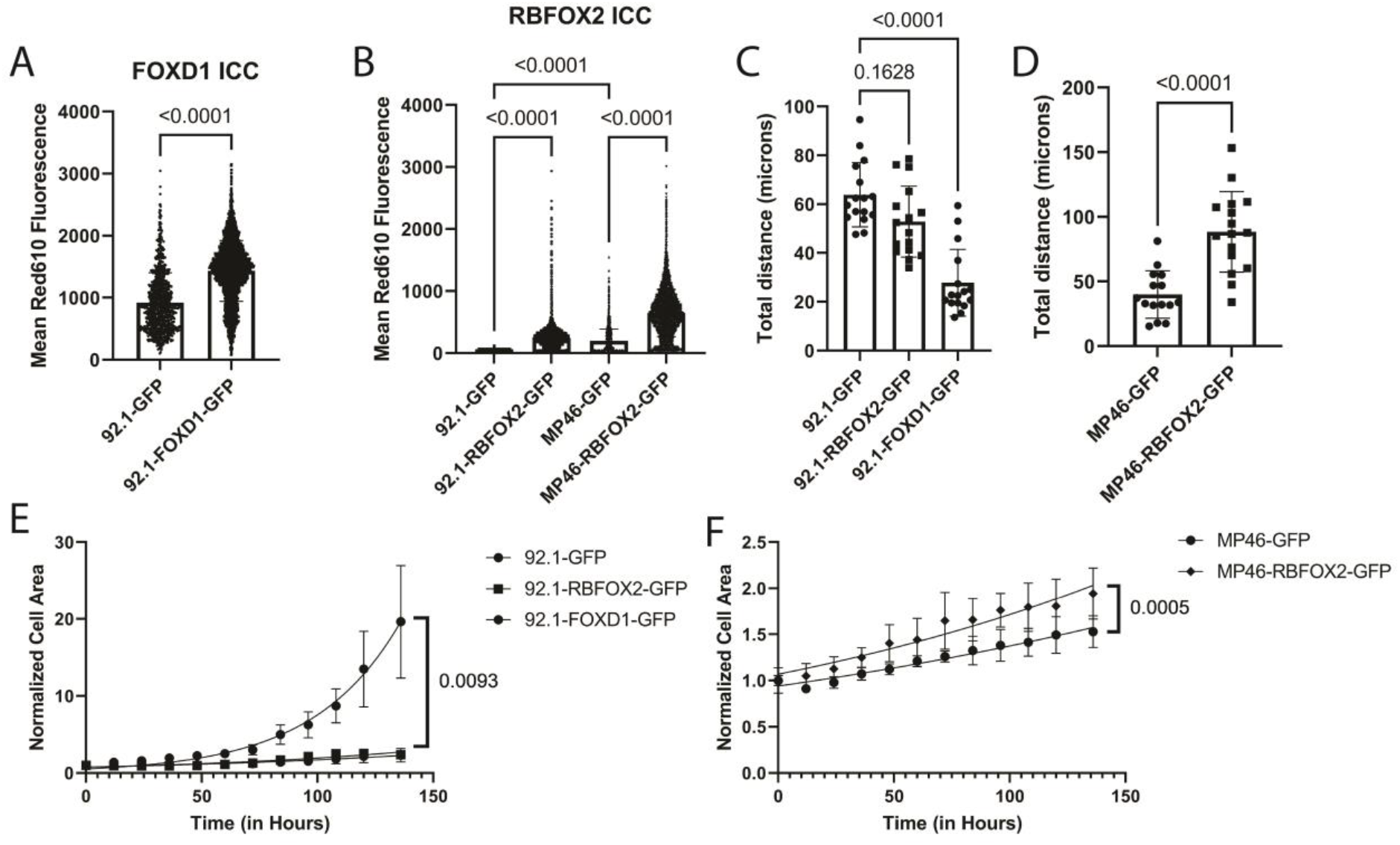
Validation and in vitro characterization of UM cell lines with overexpression of RBFOX2 or FOXD1. A) Quantification of FOXD1 protein expression using Immunocytochemistry (ICC) on 92.1-GFP and 92.1-FOXD1-GFP. B) Quantification of RBFOX1 protein expression using ICC on 92.1-GFP, 92.1-RBFOX2-GFP, MP46-GFP and MP46-RBFOX2-GFP. C-D) Total migratory distance after 24 hours using wound healing assays. E) Cell proliferation assay of 92.1-GFP, 92.1-RBFOX2-GFP and 92.1-FOXD1-GFP over a 144 hour time period. F) Cell proliferation assay of 92.1-GFP, 92.1-RBFOX2-GFP and 92.1-FOXD1-GFP over a 144 hour time period.

**Figure 5:**
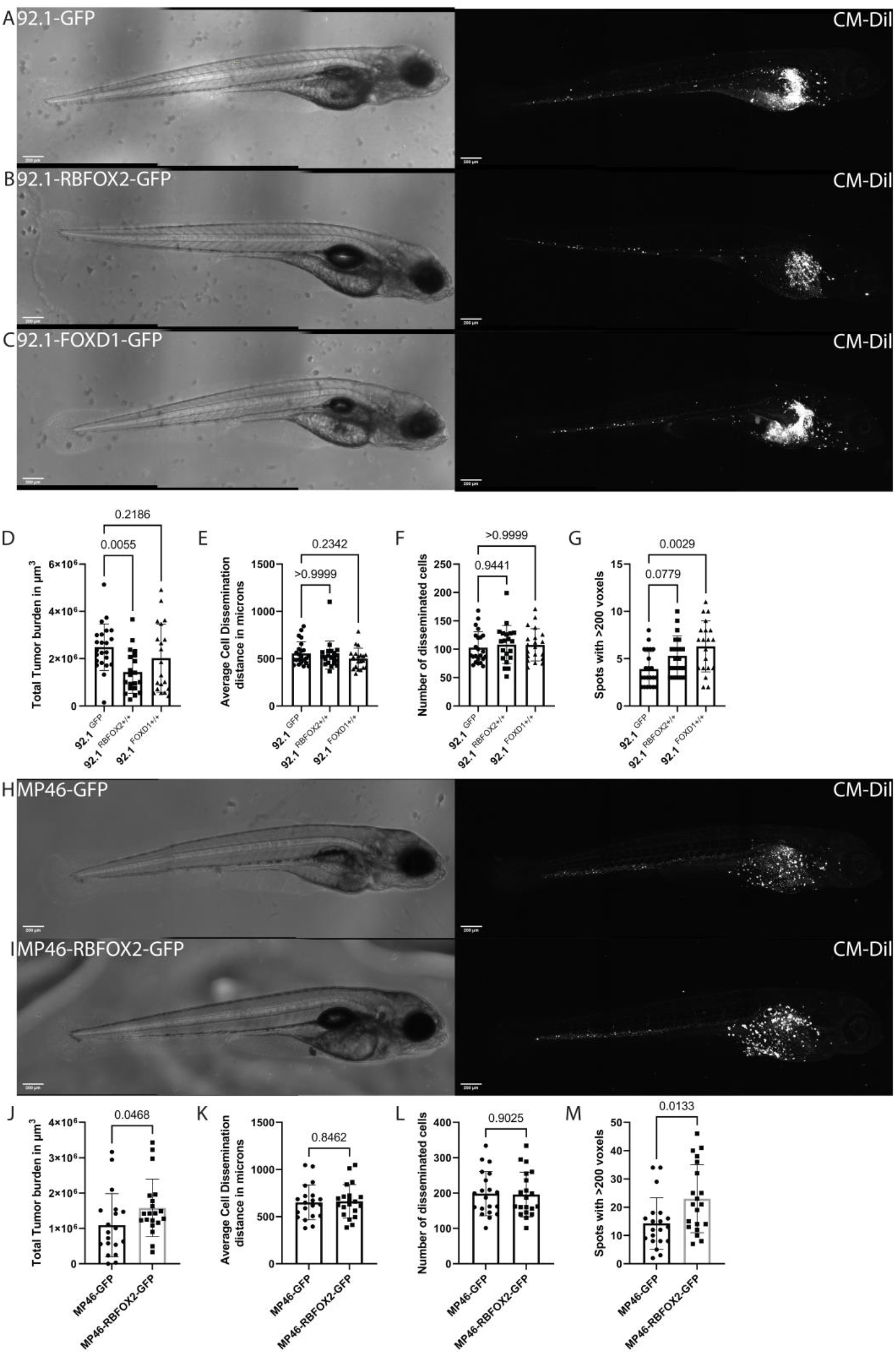
Characterization of in vivo behavior of UM cell lines containing overexpression of RBFOX2 or FOXD1. A-C) Representative images of 92.1-GFP, 92.1-RBFOX2-GFP and 92.1-FOXD1-GFP xenograft models. D) Quantification of total tumor burden in 92.1-GFP xenograft models. E) Quantification of cell dissemination distances in 92.1-GFP xenograft models. F) Quantification of amount of cell disseminations in 92.1-GFP xenograft models. G) Quantification of amount of large cell clusters in 92.1-GFP xenograft models H-I) Representative images of MP46-GFP and MP46-RBFOX2-GFP xenograft models. J) Quantification of total tumor burden in MP46-GFP xenograft models. K) Quantification of cell dissemination distances in MP46-GFP xenograft models. L) Quantification of amount of cell disseminations in MP46-GFP xenograft models. M) Quantification of number of large cell clusters in MP46-GFP xenograft models.

### Elevated RBFOX2 or FOXD1 expression improves cell survival outside of inoculation sites in zebrafish xenografts dependent on cell type

Zebrafish xenografts were used to examine whether RBFOX2 or FOXD1 enhances aggressive behavior in vivo. Interestingly, overexpression of either RBFOX2 or FOXD1 did not increase the tumor burden, number of disseminated cells or their distance from the inoculation site in 92.1-xenografts (Figure 5D-F). However, the number of large cell clusters outside of the inoculation site was increased in 92.1-FOXD1-GFP-xenografts compared to GFP controls (Figure 5G). Although RBFOX2 overexpression did not alter 92.1-cells, MP46 cells drastically changed. MP46-RBFOX2-GFP-xenografts demonstrate a significantly higher tumor burden (Figure 5J). Although the number of disseminated cells and their distance did not change (Figure 5K, 5L), the number of large cell clusters was significantly increased (Figure 5M). This suggests that RBFOX2 is able to improve cell survival and potentially cell proliferation outside of the inoculation site in a cell dependent fashion.

## Discussion

Previous identification of the clinical correlations of RBFOX2 and FOXD1 provided novel biomarkers. Yet, correlation does not mean causation. Therefore, this study investigated if and how RBFOX2 and FOXD1 play a role in uveal melanoma progression. Our results illuminate their context-dependent contribution to migratory behavior and dissemination, underscoring the complexity of RBFOX2 and FOXD1 in relation to BAP1 mutational status.

We demonstrate both *in vitro* and *in vivo that* loss of RBFOX2 inhibits both migratory and cell growth capacity with the largest effect in BAP1^neg^ cells. Interestingly, overexpression of RBFOX2 did not elevate aggressive behavior in BAP1^pos^ cells, whereas BAP1^neg^ cells did show increased aggressiveness *in vivo*. These result underscore the mutation-dependent role of RBFOX2 in uveal melanoma. RBFOX2 functions as a complex regulatory protein that governs a wide array of transcripts through interactions with the RNA at splice sites. It has been shown to directly bind RNA and indirectly through the recruitment of partner proteins, enabling alternative splice sites that contribute to the extensive heterogeneity of RBFOX2 targets and splice outcomes[20]. Pan-cancer analysis of RBFOX2 expression reflects this heterogeneity, where poor prognosis is correlated to both high- and low-expression of RBFOX2[21]. Moreover, alternative splicing via RBFOX2 has been identified as a regulator for metastatic potential in breast [22] and laryngeal cancer [23] by facilitating an unique mechanism dependent on their specific context. Additionally, RBFOX2 has been previously associated with the Hippo-YAP signaling pathway, which is recognized as a critical pathway in UM etiology. The expression of RBFOX2 may lead to the retention of exon 6 of downstream target *TEAD1*, which was shown to increase the oncogenic potential of *TEAD1*[24]. This highlights the need for further investigation of the biochemical interactions of RBFOX2 in uveal melanoma.

In a similar fashion, we elucidate the mutation-dependent role of FOXD1 in uveal melanoma cells. The loss of FOXD1 expression resulted in a significantly reduction of tumor burden *in vivo* of both BAP1^pos^ and BAP1^neg^ cells. However, migration capacity and cell dissemination was only reduced in BAP1^neg^ cells. Furthermore, our data suggests there is a threshold of FOXD1 expression due to the lethality seen in BAP1^neg^ cells upon overexpression. Conversely, BAP1^pos^ cells overexpressing FOXD1 demonstrated minimal changes, suggesting that this threshold may be context dependent. Elevated FOXD1 expression has been reported to be associated with poor prognosis in many cancer types[25]. A recurring function of FOXD1 in cancer is its role in plasticity and resistance to therapy. FOXD1 demonstrated activation of downstream target SNAI2 in oral squamous cell carcinoma, resulting in increased plasticity and EMT[26]. However, FOXD1 promoted plasticity by activating SNAI1 in esophageal squamous cell carcinoma[27]. Moreover, FOXD1 has been demonstrated to regulate aerobic glycolysis to facilitate cell proliferation, invasion and metastasis in pancreatic cancer[28]. Collectively, these findings support our hypothesis regarding the context-dependent effects of FOXD1, suggesting that in uveal melanoma this may be mutation-dependent.

In conclusion, this study emphasizes the critical roles of RBFOX2 and FOXD1 in the progression of UM, particularly in BAP1^neg^ uveal melanoma cells. Losing RBFOX2 or FOXD1 expression reduced tumor burden, migration and dissemination, while overexpression promotes aggressive behavior. Despite evident cellular alterations, the precise underlying mechanisms remain to be fully elucidated. Current ongoing RNA sequencing efforts may provide insight into the specific roles of RBFOX2 and FOXD1 in a mutation-dependent context. Additionally, given the role in alternative splicing by RBFOX2, a comprehensive analysis of isoforms is essential to understand its complex function. Ultimately, knowledge of molecular alterations due to RBFOX2 and/or FOXD1 could provide novel pathways implicated in UM progression.

## Funding

This research was funded by the Henkes foundation (2021-04) J.Q.N.N., E.K.; Rotterdamse Blindenbelangen (HV/AB/B20200043) Q.C.C.v.d.B., E.B.; Collaborative Ophthalmic Research Rotterdam (6.2.0) J.Q.N.N., E.K.; Landelijke Stichting voor Blinden en Slechtzienden (2019-7) Q.C.C.v.d.B., E.B.; and Algemene Nederlandse Vereniging ter voorkoming van Blindheid (2019-07) Q.C.C.v.d.B., E.B. Additional funding for RNA-sequencing was funded by the Trustfonds foundation.

## Notes

### Competing Interest Statement

The authors have declared no competing interest.

## References

1. Carvajal, R.D., et al., Metastatic disease from uveal melanoma: treatment options and future prospects. Br J Ophthalmol, 2017. 101(1): p. 38–44.

2. Jager, M.J., et al., Uveal melanoma. Nat Rev Dis Primers, 2020. 6(1): p. 24.

3. Drabarek, W., et al., Multi-Modality Analysis Improves Survival Prediction in Enucleated Uveal Melanoma Patients. Invest Ophthalmol Vis Sci, 2019. 60(10): p. 3595–3605.

4. Van Raamsdonk, C.D., et al., Frequent somatic mutations of GNAQ in uveal melanoma and blue naevi. Nature, 2009. 457(7229): p. 599–602.

5. Van Raamsdonk, C.D., et al., Mutations in GNA11 in Uveal Melanoma. New England Journal of Medicine, 2010. 363(23): p. 2191–2199.

6. Johansson, P., et al., Deep sequencing of uveal melanoma identifies a recurrent mutation in PLCB4. Oncotarget, 2016. 7(4): p. 4624–31.

7. Nell, R.J., et al., Involvement of mutant and wild-type CYSLTR2 in the development and progression of uveal nevi and melanoma. BMC Cancer, 2021. 21(1): p. 164.

8. Vader, M.J.C., et al., GNAQ and GNA11 mutations and downstream YAP activation in choroidal nevi. Br J Cancer, 2017. 117(6): p. 884–887.

9. van Poppelen, N.M., et al., Genetic Background of Iris Melanomas and Iris Melanocytic Tumors of Uncertain Malignant Potential. Ophthalmology, 2018. 125(6): p. 904–912.

10. Yavuzyigitoglu, S., et al., Uveal Melanomas with SF3B1 Mutations: A Distinct Subclass Associated with Late-Onset Metastases. Ophthalmology, 2016. 123(5): p. 1118–28.

11. Matatall, K.A., et al., BAP1 deficiency causes loss of melanocytic cell identity in uveal melanoma. BMC Cancer, 2013. 13: p. 371.

12. Vaquerizas, J.M., et al., A census of human transcription factors: function, expression and evolution. Nature Reviews Genetics, 2009. 10(4): p. 252–263.

13. Bushweller, J.H., Targeting transcription factors in cancer — from undruggable to reality. Nature Reviews Cancer, 2019. 19(11): p. 611–624.

14. Bradner, J.E., D. Hnisz, and R.A. Young, Transcriptional Addiction in Cancer. Cell, 2017. 168(4): p. 629–643.

15. van den Bosch, Q.C.C., et al., FOXD1 Is a Transcription Factor Important for Uveal Melanocyte Development and Associated with High-Risk Uveal Melanoma. Cancers (Basel), 2022. 14(15).

16. De Waard-Siebinga, I., et al., Establishment and characterization of an uveal-melanoma cell line. Int J Cancer, 1995. 62(2): p. 155–61.

17. Amirouchene-Angelozzi, N., et al., Establishment of novel cell lines recapitulating the genetic landscape of uveal melanoma and preclinical validation of mTOR as a therapeutic target. Mol Oncol, 2014. 8(8): p. 1508–20.

18. Brinkman, E.K., et al., Easy quantitative assessment of genome editing by sequence trace decomposition. Nucleic Acids Research, 2014. 42(22): p. e168–e168.

19. van den Bosch, Q.C.C., E. Kiliç, and E. Brosens, Uveal Melanoma Zebrafish Xenograft Models Illustrate the Mutation Status-Dependent Effect of Compound Synergism or Antagonism. Investigative Ophthalmology & Visual Science, 2024. 65(10): p. 26–26.

20. Zhou, D., et al., RBFOX2 alters splicing outcome in distinct binding modes with multiple protein partners. Nucleic Acids Research, 2021. 49(14): p. 8370–8383.

21. Huang, F., et al., Integrated pan-cancer analysis reveals the immunological and prognostic potential of RBFOX2 in human tumors. Frontiers in Pharmacology, 2024. 15.

22. Shapiro, I.M., et al., An EMT-driven alternative splicing program occurs in human breast cancer and modulates cellular phenotype. PLoS Genet, 2011. 7(8): p. e1002218.

23. Lu, B., et al., LncRNA ZFAS1 promotes laryngeal cancer progression through RBFOX2-mediated MENA alternative splicing. Environ Toxicol, 2023. 38(3): p. 522–533.

24. Choi, S., et al., RBFOX2-regulated TEAD1 alternative splicing plays a pivotal role in Hippo-YAP signaling. Nucleic Acids Res, 2022. 50(15): p. 8658–8673.

25. Cheng, L., et al., Dissecting multifunctional roles of forkhead box transcription factor D1 in cancers. Biochimica et Biophysica Acta (BBA) - Reviews on Cancer, 2023. 1878(6): p. 188986.

26. Chen, Y., et al., FOXD1 promotes EMT and cell stemness of oral squamous cell carcinoma by transcriptional activation of SNAI2. Cell & Bioscience, 2021. 11(1): p. 154.

27. Wu, Z., et al., FOXD1 promotes proliferation, migration, and epithelial-mesenchymal transformation in esophageal squamous cell carcinoma by targeting SNAI1. Process Biochemistry, 2024. 138: p. 47–56.

28. Cai, K., et al., FOXD1 facilitates pancreatic cancer cell proliferation, invasion, and metastasis by regulating GLUT1-mediated aerobic glycolysis. Cell Death & Disease, 2022. 13(9): p. 765.

