## Supplementary File for "Transcriptional regulators FOXD1 and RBFOX2 contribute to metastatic capacity in *BAP1*^*mut*^ uveal melanoma"

**Contents:**

**Supplementary figure 1:** **Validation of stable knock-out cell lines**

**Supplementary figure 2:** **Representation of cell attachment analysis of UM cell lines**

**Supplementary figure 3: Cell tracking of wild-type and knock-out UM cell lines**

**Supplementary figure 4: Plasmid Maps of pFOXD1-GFP and pRBFOX2-GFP**

**Supplementary figure 5: Immunocytochemistry of FOXD1 and RBFOX2 on established overexpression UM cell lines**

**Supplementary figure 6: Morphometric analysis of 92.1 and MP46 cells with overexpression of RBFOX2 or FOXD1.**

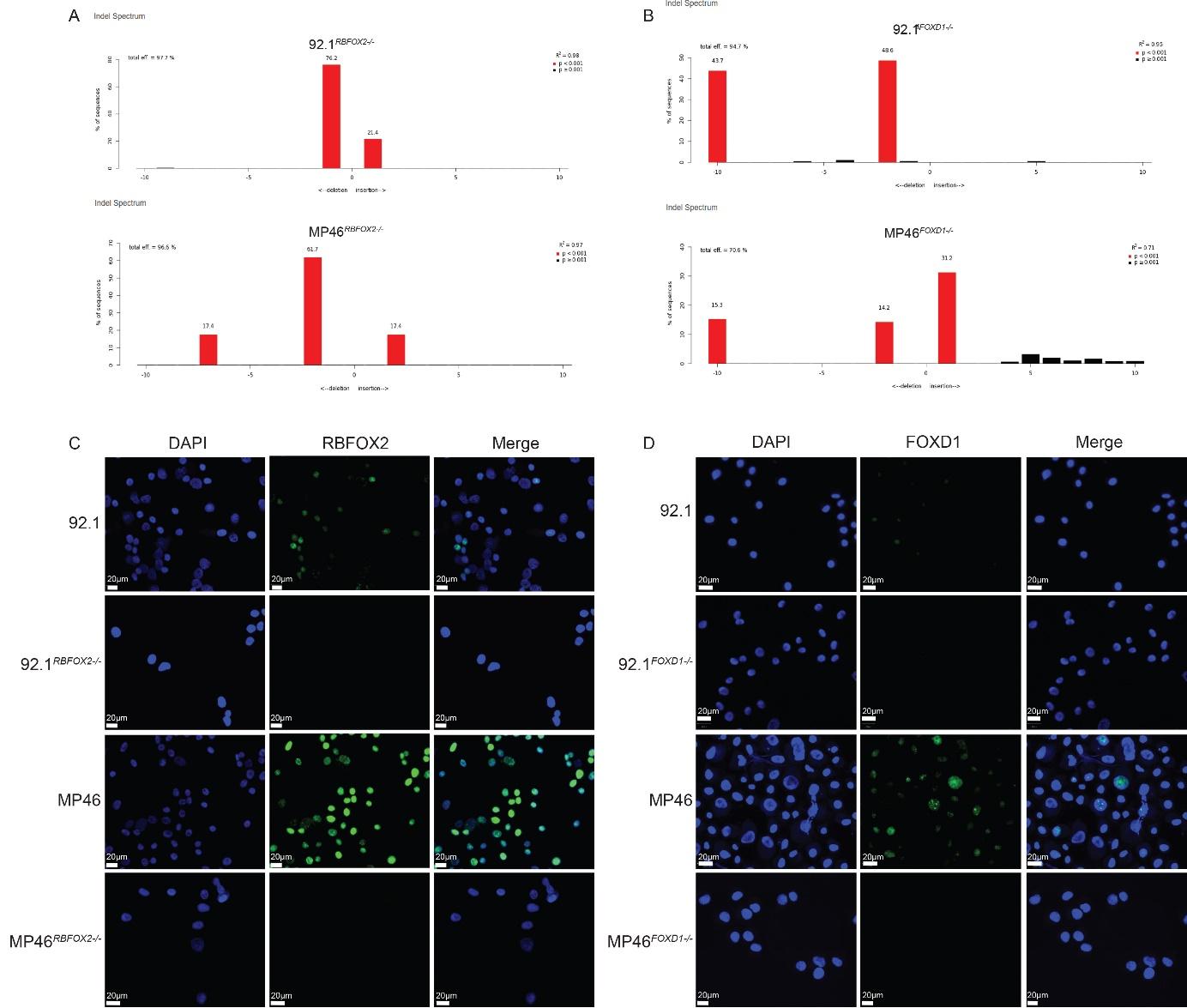

***Supplementary figure 1:Validation of stable knock-out cell lines.*** *Cell lines 92.1 (BAP1pos) and MP46 (BAP1neg) were sequenced after transfection of CRISPR-Cas9 plasmids with targeted sgRNAs. Selected clone for this study lack any wild-type alleles for RBFOX2 (S1A) or FOXD1 (S1B) as shown by TIDE analysis. Complete loss of RBFOX2 and FOXD1 was validated on RNA (Figure 1) and protein with immunocytochemistry (Figure 1, S1C,D, repsectively).*

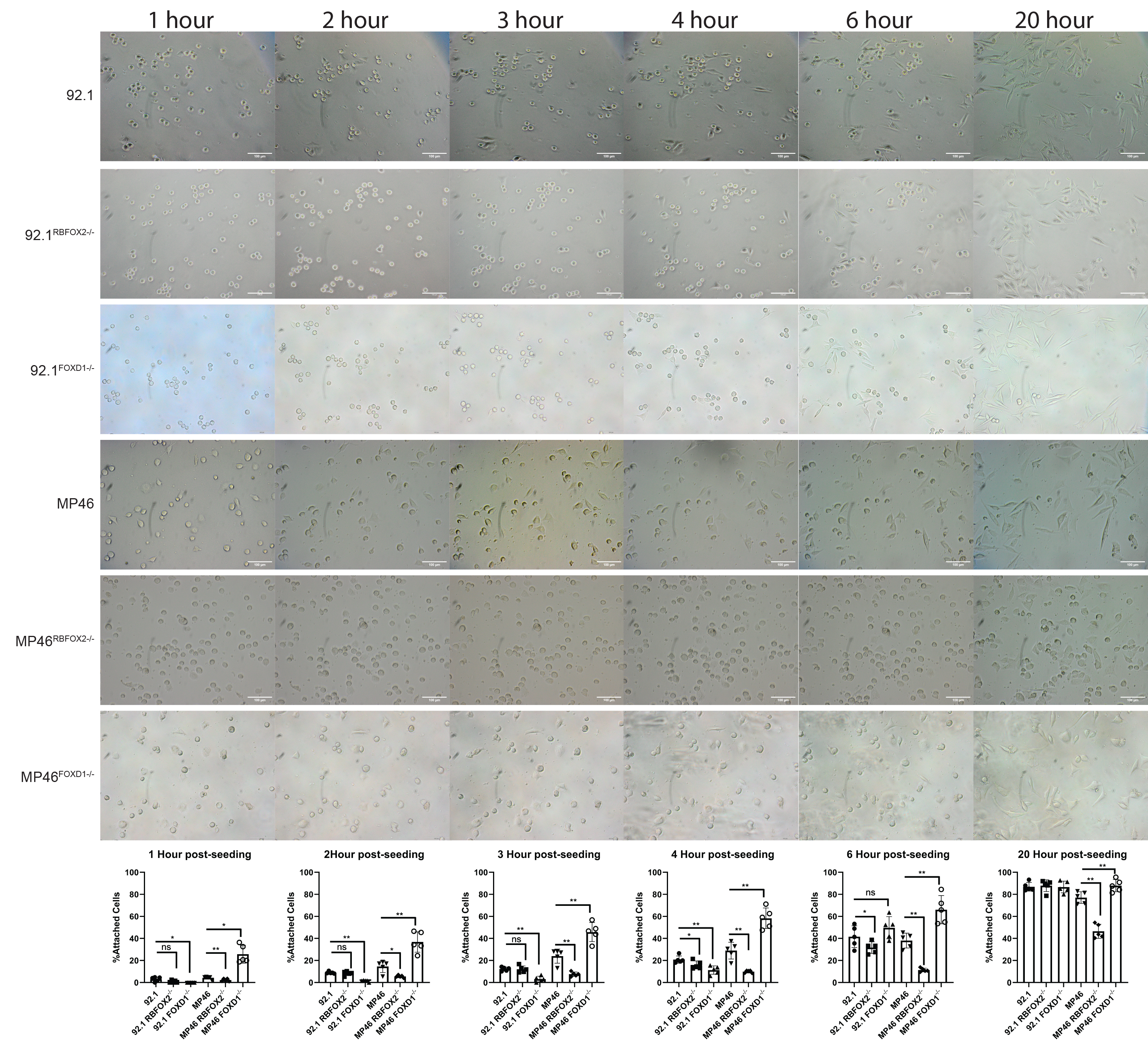

***Supplementary figure 2:Representation of cell attachment analysis of UM cell lines.*** *Representative brightfield images of cell attachment analysis of both wild/type and knock-out cell lines is shown with time-based invervals of 1, 2, 3, 4, 6 and 20 hours post tryptinization and seeding of cells. For each time point the percentage of attached cells are shown below the representative images illustrating significant differences between cell lines.*

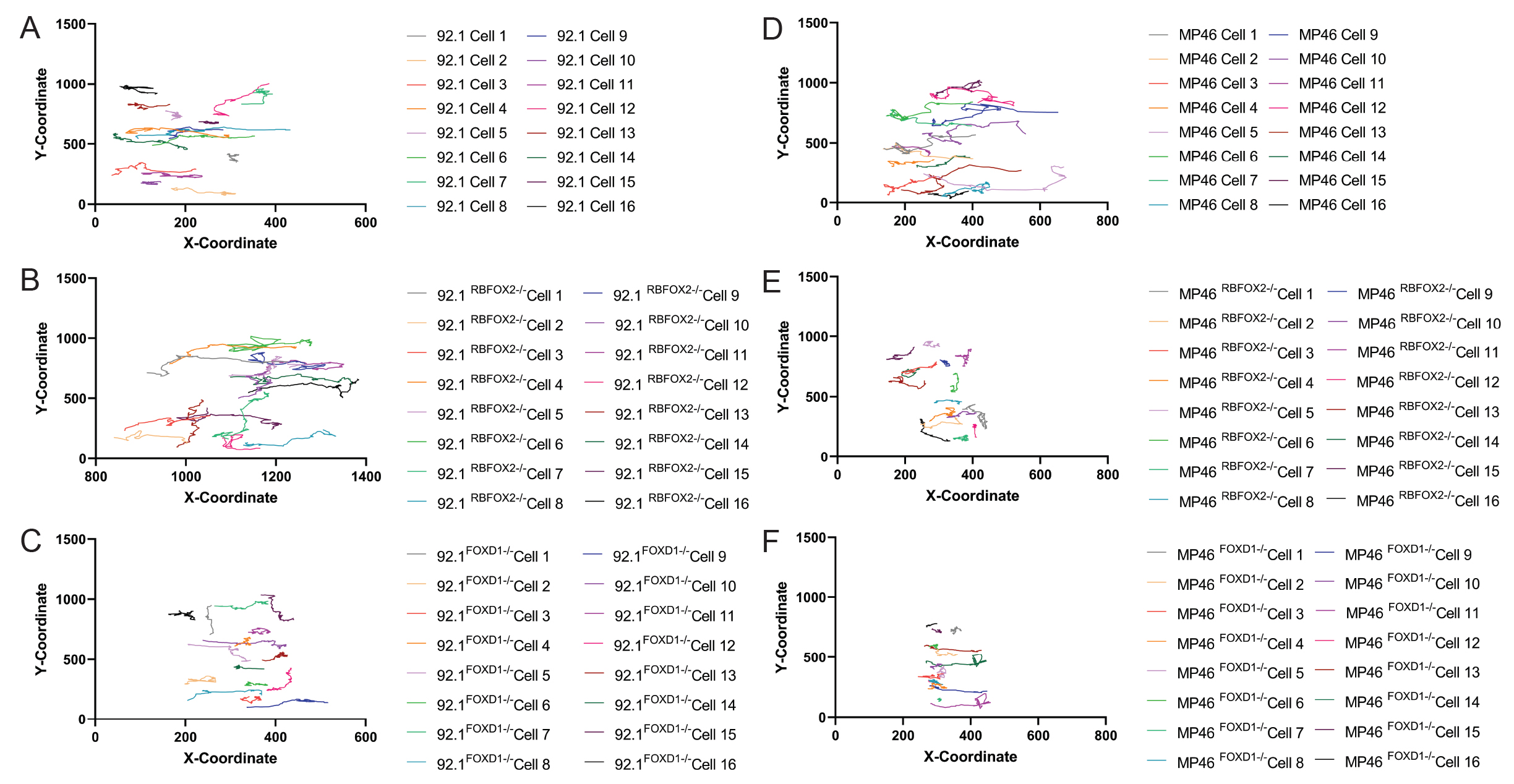

***Supplementary figure 3: Cell tracking of wild-type and knock-out UM cell lines.*** *A) migratory track of individual 92.1 cells over a time span of 24 hours. B) migratory track of individual 92.1-RBFOX2-/- cells over a time span of 24 hours. A) migratory track of individual 92.1-FOXD1-/- cells over a time span of 24 hours. A) migratory track of individual MP46 cells over a time span of 24 hours. A) migratory track of individual MP46-RBFOX2-/- cells over a time span of 24 hours. A) migratory track of individual MP46-FOXD1-/- cells over a time span of 24 hours.*

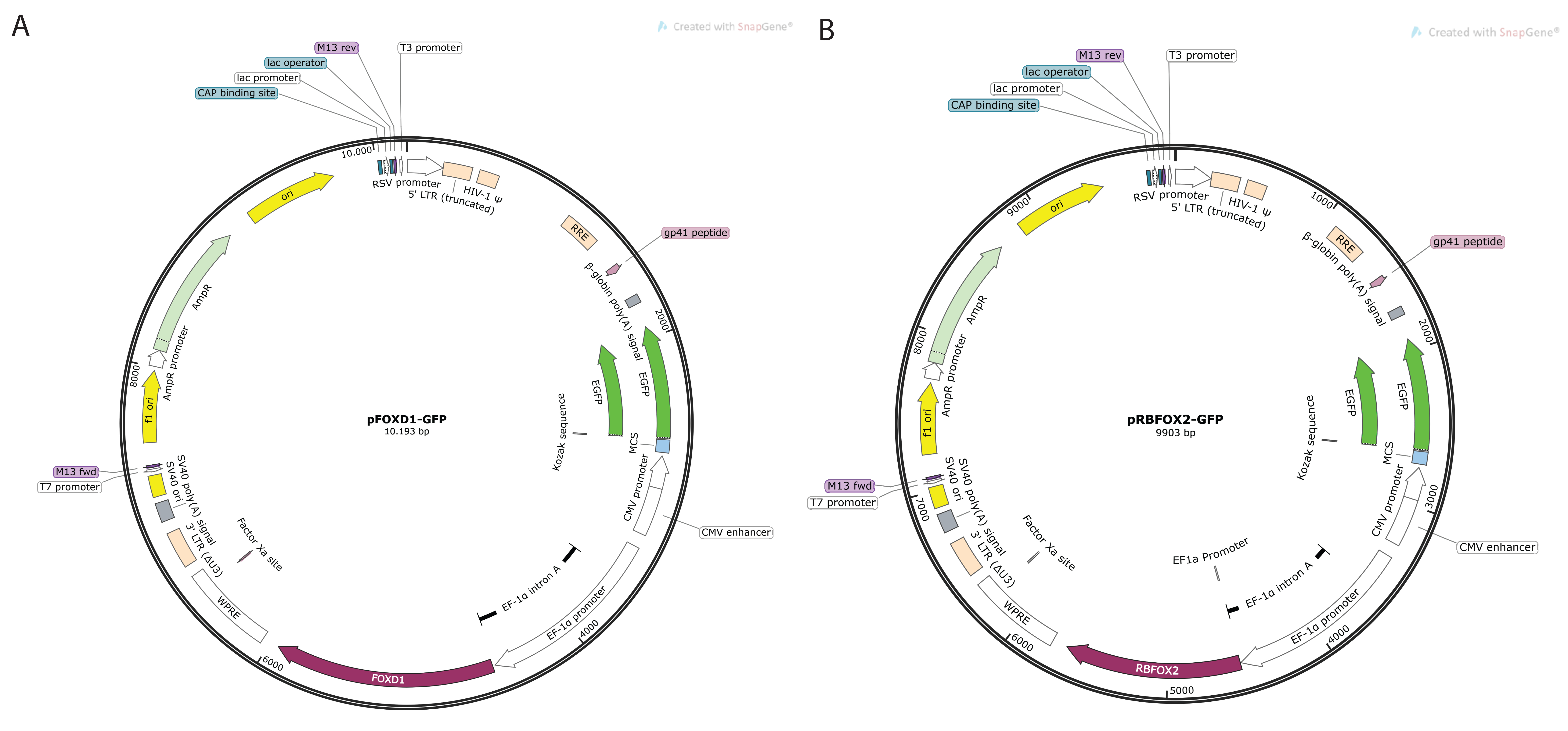

***Supplementary figure 4: Plasmid Maps of pFOXD1-GFP and pRBFOX2.*** *Adapted viral plasmids to allow for genomic integration of FOXD1 (A) or RBFOX2 (B), with GFP as a selection marker.*

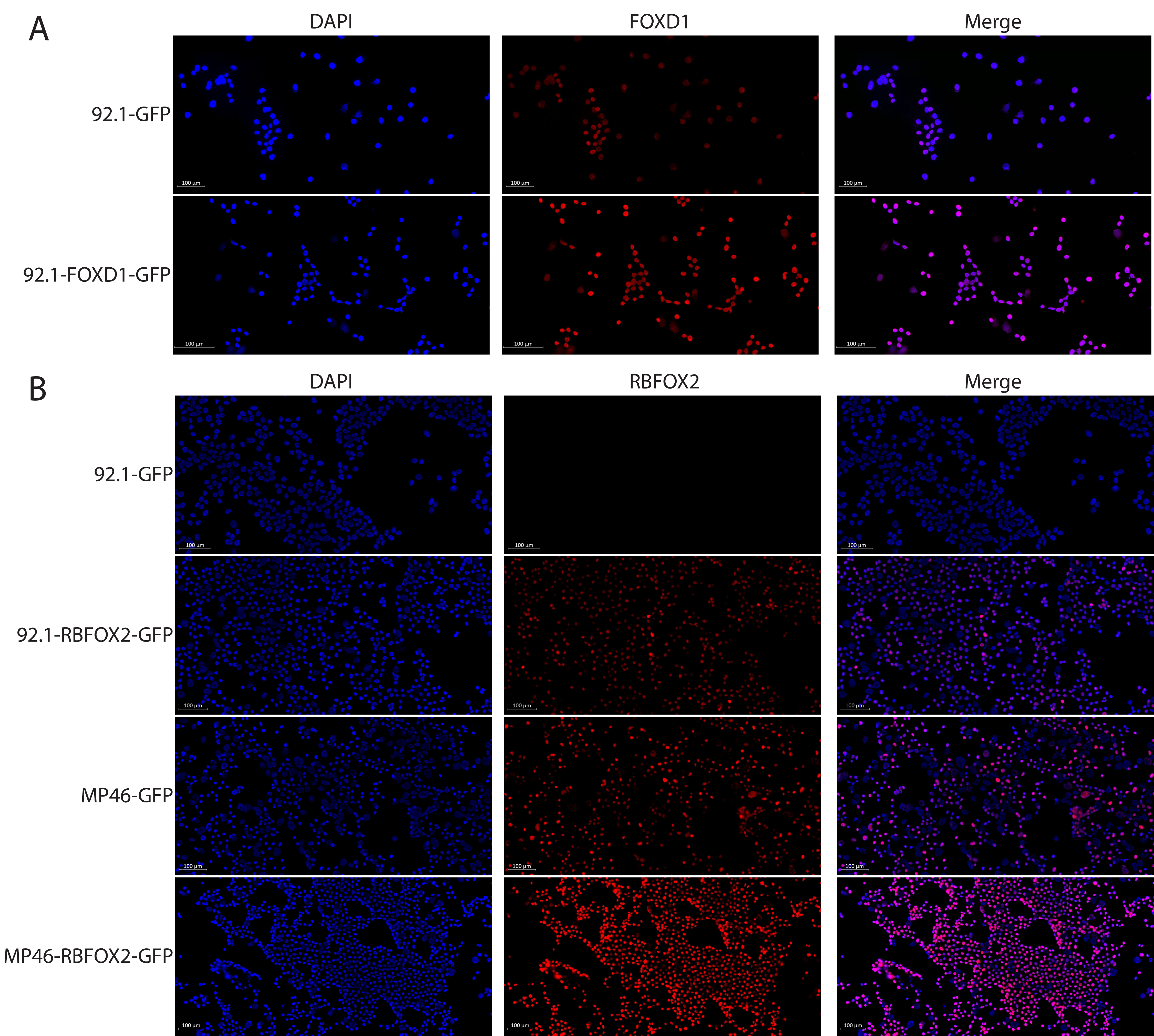

***Supplementary figure 5: Immunocytochemistry of FOXD1 and RBFOX2 on established overexpression UM cell lines.*** *A) FOXD1 immunocytochemistry on 92.1-GFP and 92.1-FOXD1-GFP. B) RBFOX2 immunocytochemistry on 92.1-GFP, 92.1-RBFOX2-GFP, MP46-GFP and MP46-RBFOX2-GFP. Scale bar is 100 micron.*

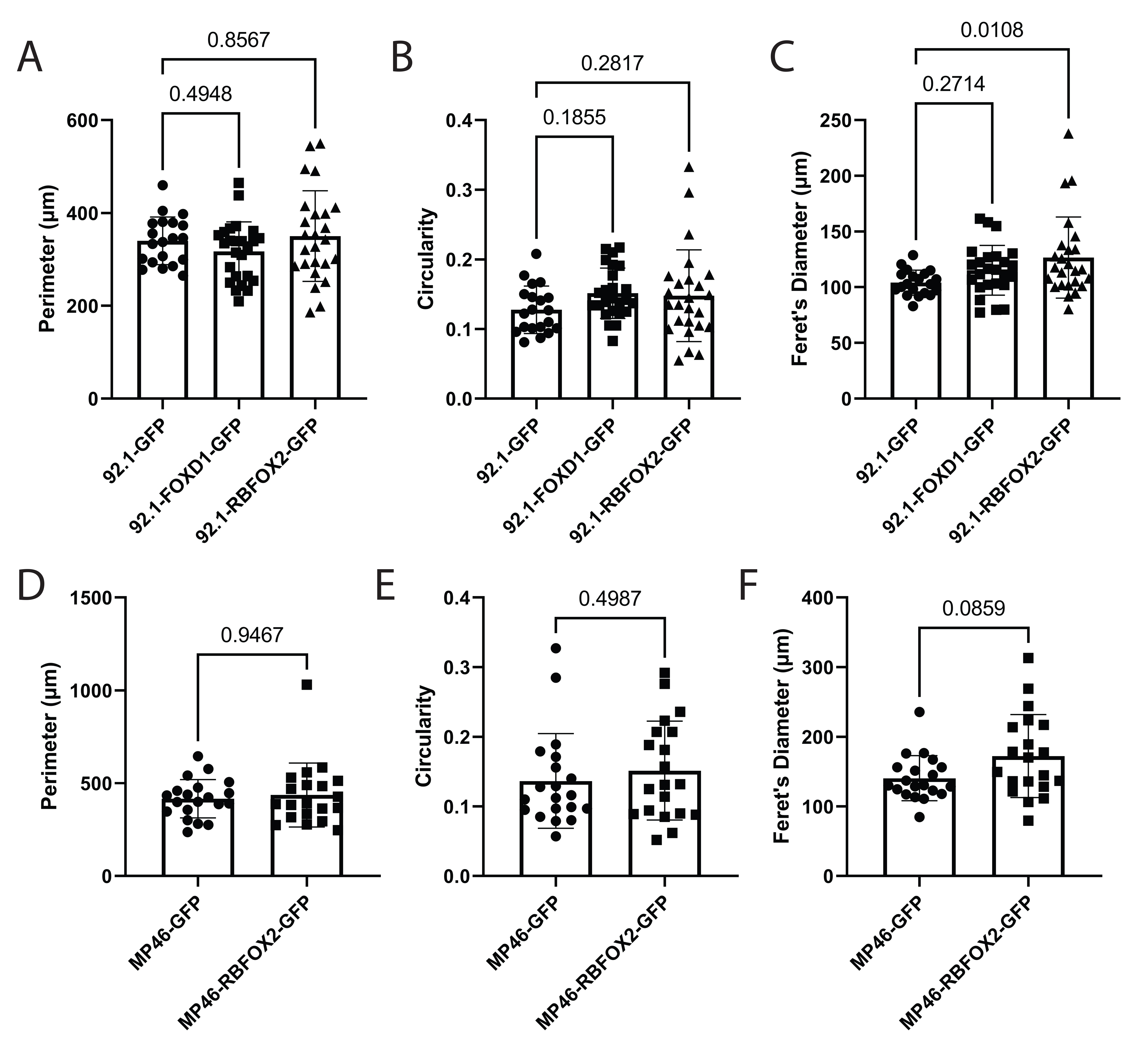

***Supplementary figure 6: Morphometric analysis of 92.1 and MP46 cells with overexpression of RBFOX2 or FOXD1.*** *A) Quantifications of perimeter size of 92.1-GFP, 92.1-FOXD1-GFP and 92.1-RBFOX2-GFP cells. B) Quantifications of the circularity of 92.1-GFP, 92.1-FOXD1-GFP and 92.1-RBFOX2-GFP cells. C) Quantifications of Feret’s diameter of 92.1-GFP, 92.1-FOXD1-GFP and 92.1-RBFOX2-GFP cells. D) Quantifications of perimeter size of MP46-GFP, and MP46-RBFOX2-GFP cells. D) Quantifications of the circularity of MP46-GFP and MP46-RBFOX2-GFP cells. F) Quantifications of perimeter size of MP46-GFP and MP46-RBFOX2-GFP cells.*
